# Biogenesis of a mitochondrial DNA inheritance machinery in the mitochondrial outer membrane

**DOI:** 10.1101/190751

**Authors:** Sandro Käser, Mathilde Willemin, Felix Schnarwiler, Bernd Schimanski, Daniel Poveda-Huertes, Silke Oeljeklaus, Bettina Warscheid, Chris Meisinger, André Schneider

## Abstract

Mitochondria cannot form de novo but require mechanisms that mediate their inheritance to daughter cells. The parasitic protozoan *Trypanosoma brucei* has a single mitochondrion with a single-unit genome that is physically connected across the mitochondrial membranes to the basal body of the flagellum. This connection, termed tripartite attachment complex (TAC), is essential for the segregation of the replicated mitochondrial genomes prior to cytokinesis. Here we identify a protein complex consisting of three integral mitochondrial outer membrane proteins - TAC60, TAC42 and TAC40 - which are essential subunits of the TAC. TAC60 contains separable mitochondrial import and TAC-sorting signals and its biogenesis depends on the main outer membrane protein translocase. TAC40 is a member of the mitochondrial porin family, whereas TAC42 represents a novel class of mitochondrial outer membrane β-barrel proteins. Consequently TAC40 and TAC42 contain C-terminal β-signals. Thus in trypanosomes the highly conserved β-barrel protein assembly machinery plays a major role in the biogenesis of its unique mitochondrial genome segregation system.

## Introduction

Mitochondria are a hallmark of eukaryotic cells (1). They derive from an endosymbiotic event between an archaeal host cell and an α-proteobacterium. The bacterial symbiont was subsequently converted into an organelle. Continued evolution since the origin of the mitochondrion, approximately 1.5-2 billion years ago, has led to a great diversification of the organelle (2, 3). This is illustrated by the immense variation of the morphology and the behaviour of mitochondria in different species and the large variation in the organization and coding content of their genomes. However, faithful transmission of mitochondria and their genomes to their daughter cells is a problem essentially all eukaryotic cells need to solve (4, 5).

In contrast to most other eukaryotes trypanosomes and its relatives have a single mitochondrion that only contains a single unit mitochondrial genome, termed kinetoplast DNA (kDNA). The kDNA consists of two genetic elements the maxi- and the minicircles. The maxicircles are present in 25-50 copies and are 22 kb in length (6).

They contain a number of protein-coding genes expected to be present in the mitochondrial genome. Most of them are cryptogenes whose primary transcripts have to be edited by multiple uridine insertions and or deletions to become functional mRNAs. The minicircles are heterogenous in sequence, occur in several thousand copies and encode the guide RNAs that provide the information for RNA editing (7, 8). Maxi- and minicircle are highly topologically interlocked and build a large disc-shaped network that is physically linked to the basal body of the single flagellum via a structure termed tripartite attachment complex (TAC). The three zones of the TAC include the unilateral filaments that connect the kDNA network to the inside of the inner membrane (IM), a segment containing tightly apposed detergent-resistant differentiated IM and outer membranes (OM) and finally the exclusion zone filaments that build a bridge from the OM to the basal body (9-11).

Due to the single unit nature of the kDNA network, its replication and segregation need to be tightly coordinated. kDNA replication occurs at a precise stage of the cell cycle immediately before the onset of the S phase in the nucleus (12, 13). During replication the kDNA doubles in size forming a dumbbell-shaped network. The process has been studied in detail and involves numerous protein factors. The segregation of the replicated kDNA networks depends on an intact TAC. Thus during kDNA replication a new TAC forms that connects the basal body of the new flagella with the replicating kDNA network. Division of the replicated kDNA network finally is linked to the segregation of the old and the new basal bodies. Initially the two kDNA networks remain connected by a structure termed "nabelschnur" which becomes resolved when the distance between the two networks is larger than 1 μΜ. The segregation then continues and the basal bodies with the attached kDNA networks move further apart (14, 15). Subsequently, prior to cytokinesis, the mitochondrion is divided in two the plane of division intersecting between the two kDNA networks (16).

Presently only a few components of the TAC have been identified. The first one was p166 (17): it contains a single predicted transmembrane domain (TMD) which however is not essential for its localization making it difficult to decide whether p166 is a TAC protein of the IM or the unilateral filaments. TAC102 is a component of the unlilateral filaments (18), whereas p196 (19) and TAC65 (20) were shown to be extracellular TAC subunits that localize to the exclusion zone filaments. The same was the case for an as yet unkown antigen recognized by the monoclonal antibody Mab 22 (21). Furthermore, TAC40 and peripheral archaic translocase of the <u>OM</u> 36 (pATOM36), two OM proteins that localize to the TAC, have also been described (20, 22). TAC40 is a β-barrel protein of the mitochondrial porin family. pATOM36 is unusual as it is localized in the TAC region but also present all over the OM. The dual localization of pATOM36 reflects its dual function in kDNA inheritance and in the biogenesis of a subset of α-helically anchored OM proteins including most subunits of the archaic translocase of the <u>OM</u> (ATOM) (20, 23). Complementation experiments revealed that the two functions are distinct, since the C-terminus is only essential for the biogenesis of OM proteins but not for the segregation of the kDNA. Thus, pATOM36 is an integrator of mitochondrial protein import and mitochondrial genome inheritance (20, 24).

Unlike other OM proteins, the OM subunits of the TAC not only need to be targeted to mitochondria and inserted into OM, but they also have to be sorted to the region of the OM close to the single unit kDNA, where a new TAC is being formed.

Here we have discovered two novel subunits of the TAC that are integral mitochondrial OM proteins. Moreover we provide a detailed analysis of the biogenesis pathways of the two newly discovered TAC subunits as well as of the previously characterized TAC40.

We show that one of the new subunits contains separable targeting signals for import into the organelle and for sorting to the TAC, whereas the other one defines a novel class of mitochondrial β-barrel proteins.

## Results

### TAC40, TAC60 and TAC42 form a complex

Recently we have shown that the mitochondrial β-barrel protein TAC40 exclusively localizes to the TAC and is essential for its function (22). In order to identify further TAC subunits we performed SILAC-based immunoprecipitation (IP) experiments using 29-13 *T. brucei* cells and a cell line expressing in situ HA-tagged TAC40. The two cell lines were grown in the presence of isotopically-labeled heavy or light lysine and arginine. Subsequently, identical cell numbers from both populations were mixed and whole cell lysates were prepared which were subjected to IP using anti-HA antibodies. The resulting eluates were analyzed by quantitative MS and Fig. 1A shows that they contain only two proteins that were five-fold or more enriched. One was TAC40, whose tagged variant served as a bait, the other one a 60 kDa protein, termed TAC60 (Tb927.7.1400). Subsequently a cell line allowing tetracycline-inducible expression of a Myc-tagged version of TAC60 was used to do a second set of SILAC-IPs. The most highly enriched proteins recovered in these IPs were TAC60, TAC40 and a protein of 42 kDa, termed TAC42 (Fig. 1B). Finally we produced a cell line expressing an HA-tagged version of TAC42 and performed a third set of SILAC-IPs which besides TAC42 itself recovered TAC40 and TAC60 as the most enriched proteins (Fig. 1C). In summary these reciprocal IPs show that TAC40, TAC60 and TAC42 form a protein complex.

**Figure 1.**
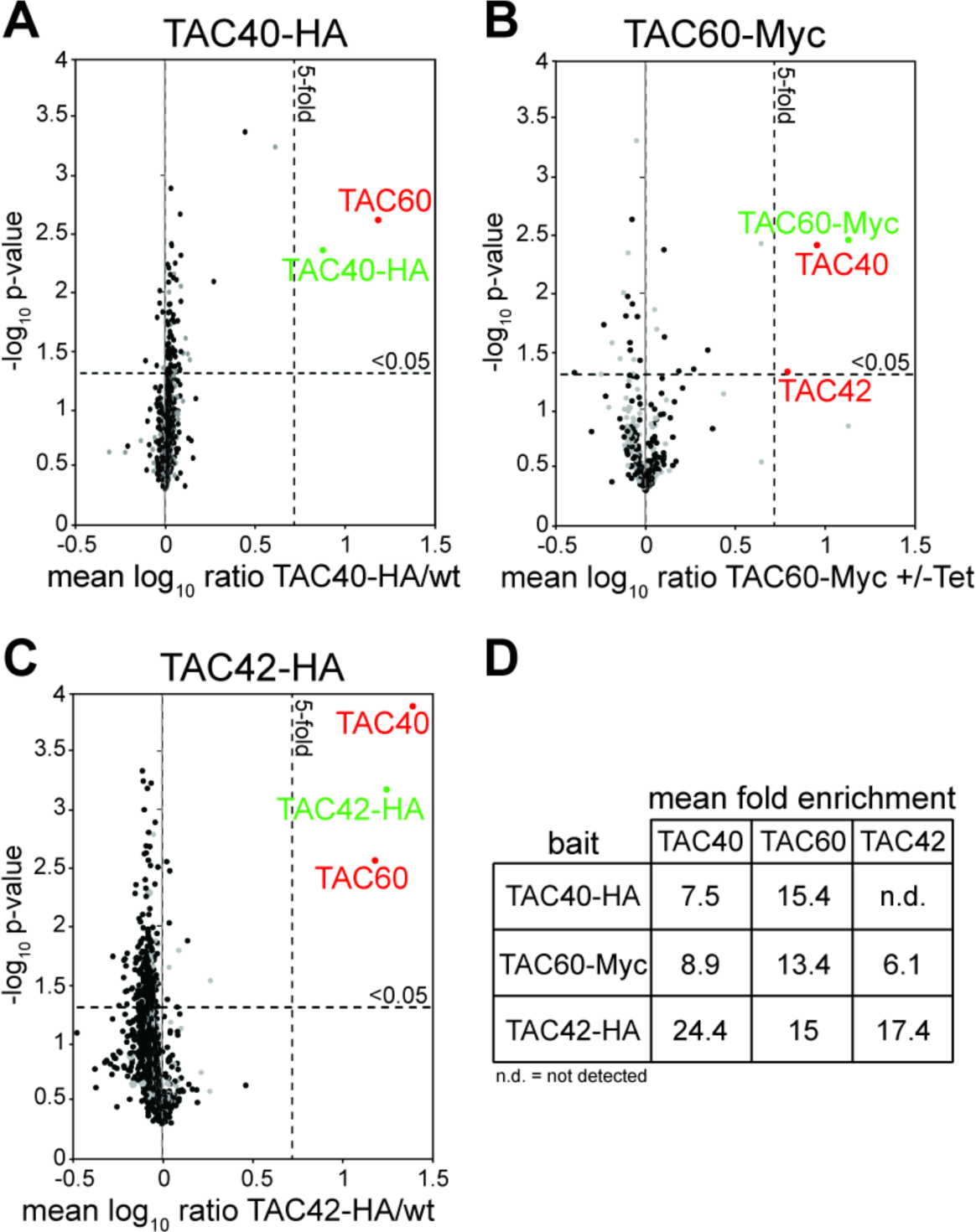
TAC40, TAC60 and TAC42 form a complex. (A) SILAC-IP of C-terminally HA-tagged TAC40 from digitonin-solubilized whole cell lysates. Mean log10 ratios (TAC40-HA/wt) of proteins detected by quantitative MS in at least two of three independent biological replicates are plotted against the corresponding log10 *P* values (one-sided t-test). Horizontal dashed line indicates a t-test significance level of 0.05, while vertical dashed lines mark a fivefold enrichment. The bait protein TAC40 is marked in green. The co-precipitated proteins are marked in red. (B) and (C) Cell lines expressing Myc- and HA-tagged versions of the TAC40 interactors, TAC60-Myc and TAC42-HA, respectively, were used for reciprocal SILAC-IPs. For complete lists of proteins for all three IPs, see Table S1. S3. (D) Table indicating the enrichment factors of TAC40, TAC60 and TAC42 in the reciprocal IPs.

### TAC60 and TAC42 are essential TAC subunits

To find out where TAC60 is localized the cell line expressing Myc-tagged TAC60 was analyzed by immunofluorescence (IF). The cells were stained with an anti Myc-antibody, with the DNA-staining agent DAPI, which labels both the nuclei and the kDNA networks, and with the monoclonal antibody YL1/2. The latter recognizes tyrosinated α-tubulin and in *T. brucei* stains the basal bodies as well as the distal part of the subpellicular array of microtubules (25). An overlay of all three signals reveals that TAC60 localizes to a dot-like structure between the basal bodies and the kDNA networks (Fig. 2A, left panel), as would be expected for a TAC subunit. The TAC is to a large part detergent-resistant, which allows the isolation of a fraction containing flagella whose basal bodies are still connected to the kDNA (9). IF analysis shows that in such fractions Myc-tagged TAC60 localizes between the kDNA and the flagellum (Fig. 2A, right panel) indicating that the protein is a structural subunit of the TAC.

**Figure 2.**
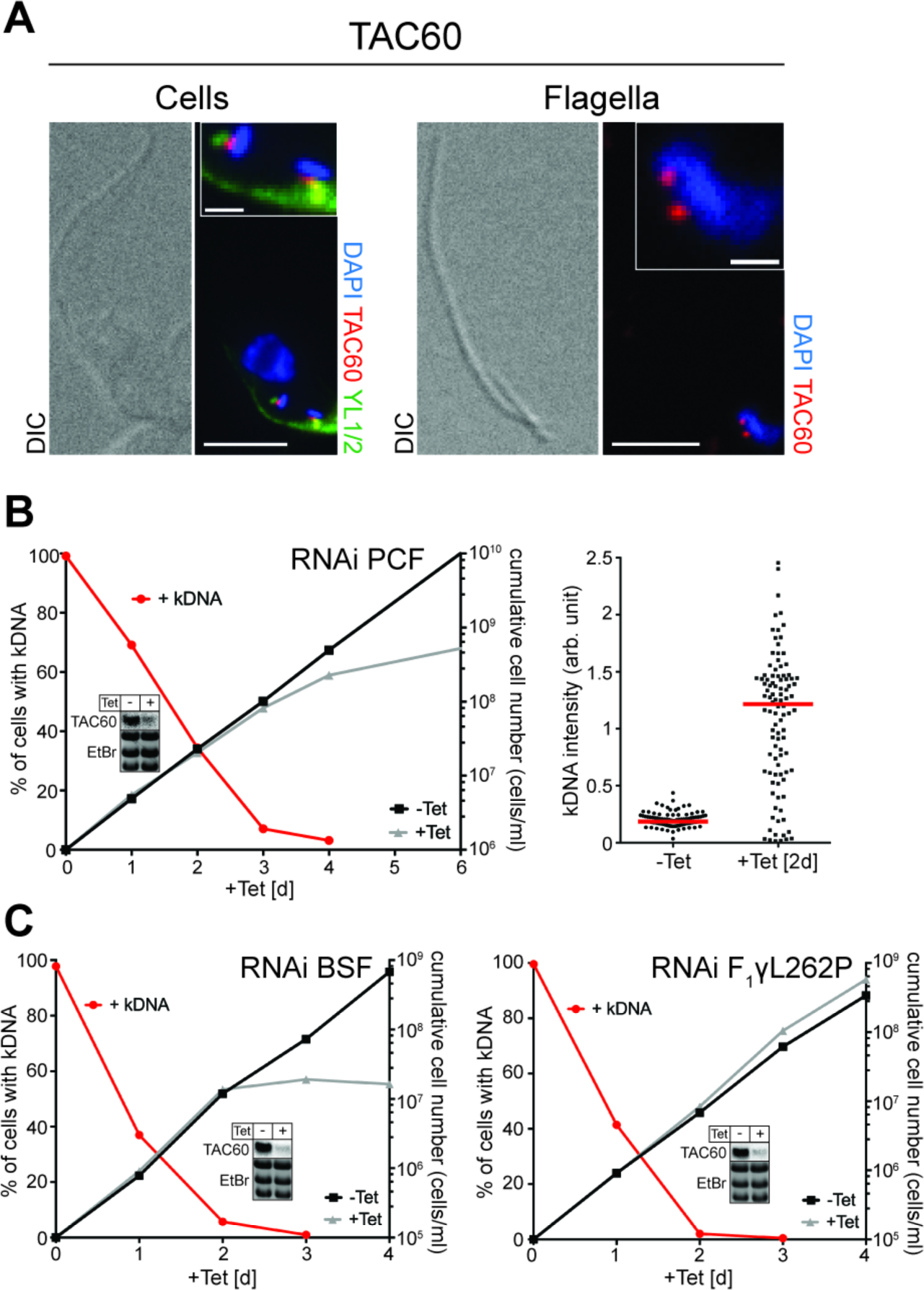
TAC60 is an essential TAC subunit. (A) Left panels: IF analysis of procyclic *T. brucei* cells expressing Myc-tagged TAC60 (red). DNA is stained with DAPI (blue). YL1/2 which serves as a marker for the basal body is indicated in green. Bar, 5 μm. Inset: magnification of the kDNA region. Bar inset, 1 μm. Right panels: IF analysis of isolated flagella of *T. brucei* cells expressing Myc-tagged TAC60. (B) Left graph: growth and loss of kDNA of the procyclic TAC60-RNAi cell line. Red lines depict percentage of cells still having the kDNA. Right graph: fluorescent intensities of kDNA networks were measured after TAC60 knock down. Red lines mark the median. (C) Left graph: growth and loss of kDNA in the bloodstream form TAC60-RNAi cell line. Right graph: growth and loss of kDNA of a TAC60-RNAi cell line from a bloodstream form *T. brucei* strain that contains a compensatory nuclear mutation (ATPase γ, L262P) that allows it to grow in the absence of kDNA (bloodstream-L262P) (27). Insets: Northern blots confirming ablation of the TAC60 mRNA in the different RNAi cell lines. Ethidium bromide-stained gel showing the rRNA region is used as a loading control.

In order to investigate the function of TAC60 a tetracycline-inducible RNAi cell line was produced. Analysis of DAPI-stained cells in the left panel of Fig. 2A shows that ablation of TAC60 in insect stage *T. brucei* leads to a rapid loss of the kDNA networks reaching approximately 50% after 1.5 days of induction. Growth on the other hand is affected after 3 days only (Fig. 2B). The right panel in Fig. 2B demonstrates that a large majority of the 30% of cells, that have retained their kDNA after two days of RNAi induction, have greatly enlarged kDNA networks (Fig. 2B, right panel). Moreover, in a minority of cells smaller kDNA networks are observed. The massive over-replication of kDNA networks is a hallmark that distinguishes cells with a deficient TAC from cells ablated in kDNA replication (17, 18, 20, 22).

Mitochondrial translation and thus the kDNA as well as the TAC are essential in both the insect and bloodstream form of *T. brucei* (26). In line with this ablation of TAC60 in bloodstream form cells leads to a rapid loss of the kDNA with a subsequent growth arrest (Fig. 2C, left panel). However, if the same experiment is done in a bloodstream form cell line of *T. brucei* that, due to a single point mutation in the nuclear-encoded γ-subunit of the mitochondrial ATPase, can grow in the absence of the kDNA (27), a different result is obtained: in such a cell line TAC function is dispensable and ablation of TAC60, while causing the loss of the kDNA, does not slow down growth (Fig. 2C, right panel).

The set of experiments done for TAC60 (Fig. 2) were also used to analyze TAC42 (Fig. 3). The obtained results were essentially identical for both proteins. The only difference was that the over-replication of the kDNA upon ablation of TAC42 was less extensive than for TAC60. In summary, these experiments (Fig. 2 and Fig. 3) establish that TAC60 and TAC42 are essential novel subunits of the TAC that are not involved in any other essential functions unrelated to the kDNA.

**Figure 3.**
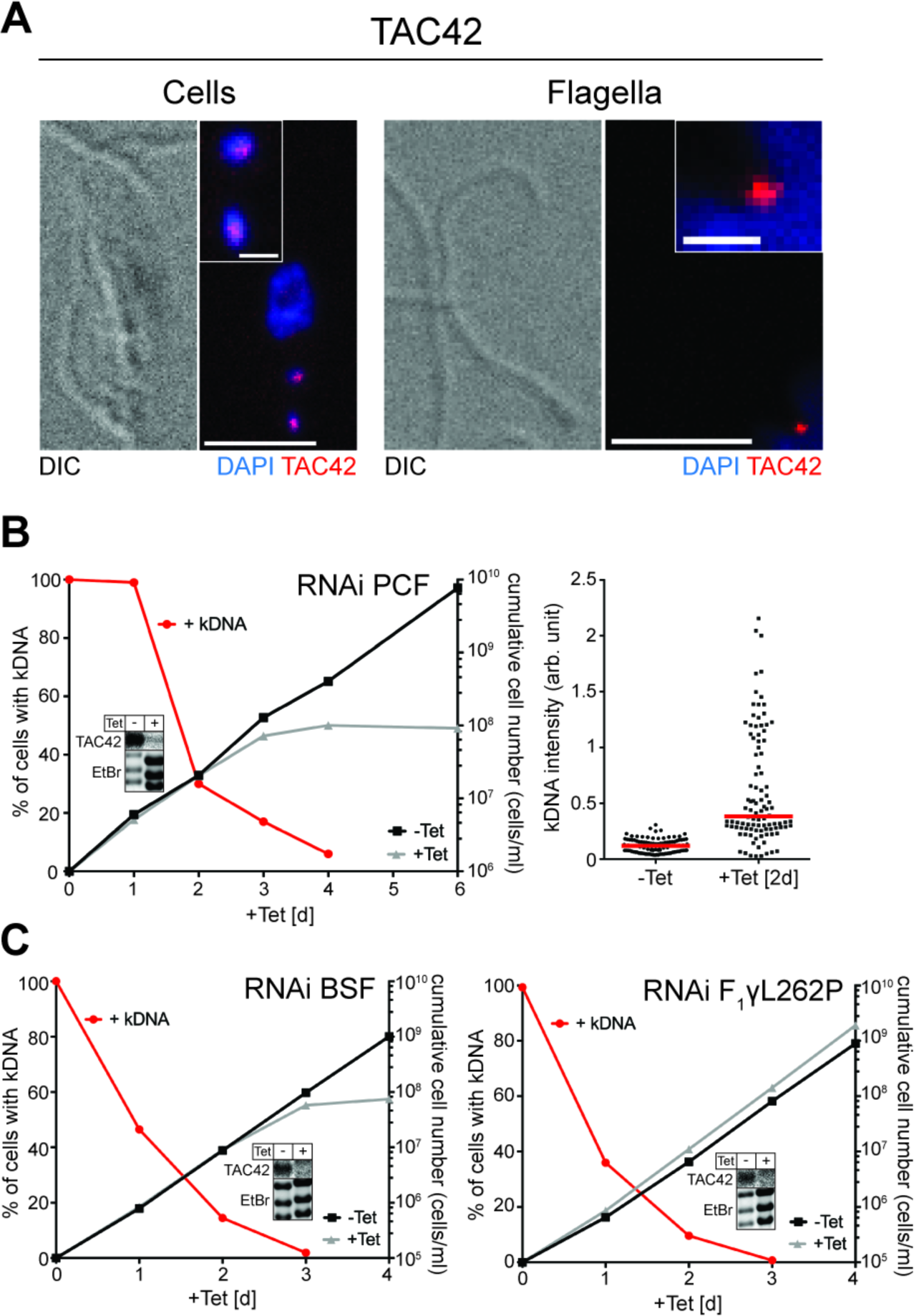
TAC42 is an essential TAC subunit. (A) Left panels: IF analysis of procyclic *T. brucei* cells expressing HA-tagged TAC42 (red). DNA is stained with DAPI (blue). Bar, 5 μm. Inset: magnification of the kDNA region. Bar inset, 1 μm. Right panels: IF analysis of isolated flagella of *T. brucei* cells expressing HA-tagged TAC42. (B) Left graph: growth and loss of kDNA of the procyclic TAC42-RNAi cell line. Red lines depict percentage of cells still having the kDNA. Right graph: fluorescent intensities of kDNA networks were measured after TAC42 knock down. Red lines mark the median. (C) Left graph: growth and loss of kDNA in the bloodstream form TAC42-RNAi cell line. Right graph: growth and loss of kDNA of a TAC42-RNAi cell line from a bloodstream form *T. brucei* strain that contains a compensatory nuclear mutation (ATPase γ, L262P) that allows it to grow in the absence of kDNA (bloodstream-L262P) (27). Insets: Northern blots confirming ablation of the TAC42 mRNA in the different RNAi cell lines. Ethidium bromide-stained gel showing the rRNA region is used as a loading control.

### TAC60 is a mitochondrial OM protein

As expected tagged TAC60 co-fractionates with the mitochondrial marker ATOM40 when cells are extracted with a low concentration of digitonin (Fig. 4A, upper panel). Moreover, TAC60 is exclusively recovered in the pellet when a crude mitochondrial fraction is subjected to carbonate extraction at high pH indicating that TAC60 is an integral membrane protein (Fig 4A, middle panel).

**Figure 4.**
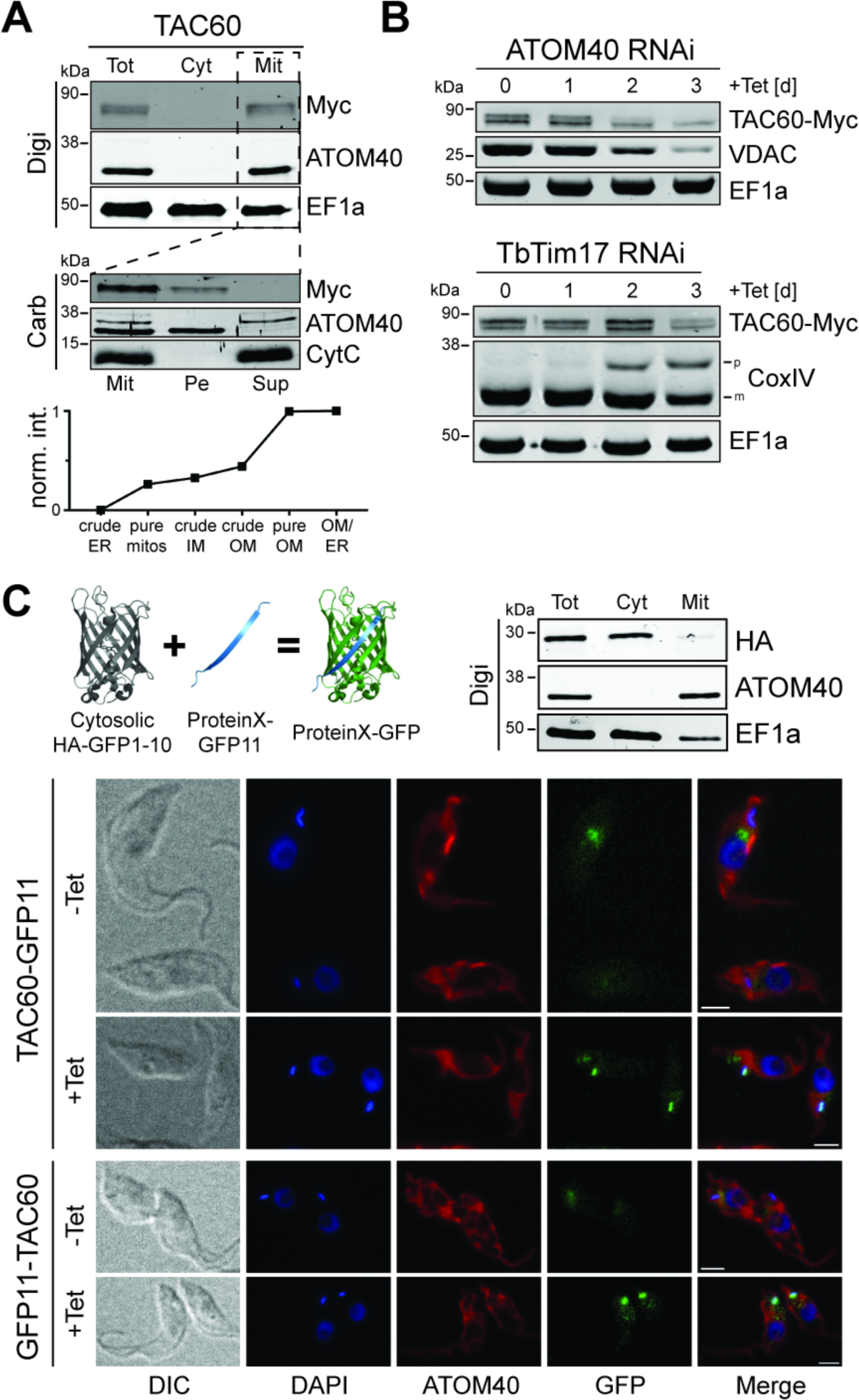
TAC60 is an OM protein whose N- and C-termini face the cytosol. (A) Top panel: immunoblot analysis of whole cells (Tot), soluble (Cyt) and digitonin-extracted mitochondria-enriched pellet (Mit) fractions of cells expressing C-terminally Myc-tagged TAC60. ATOM40 and EF1a served as mitochondrial and cytosolic markers, respectively. Middle panel: carbonate extraction at ph 11.5 of the mitochondria-enriched pellet fraction (Mit). The pellet (Pe) and the supernatant (Sup) fractions correspond integral membrane and soluble proteins. ATOM40 and cytochrome C (Cyt C) serve as markers for integral and peripheral membrane proteins, respectively. Bottom panel: normalized abundance profile of TAC60 over six subcellular fractions, produced in a previous proteomic analysis (35). (B) Immunoblots of total cellular extracts from procyclic ATOM40-RNAi (top panel) and TbTim17-RNAi (bottom panel) cells that constitutively express Myc-tagged TAC60. Time of induction is indicated. The OM protein VDAC, the IM protein CoxIV and cytosolic EF1a serve as controls. The positions of the precursor (p) and mature forms (m) of CoxIV are indicated. (C) Top left: conceptual depiction of the split GFP approach (59). Top right: immunoblot analysis of whole cells (Tot), soluble (Cyt) and digitonin-extracted mitochondria-enriched pellet (Mit) fractions of cells expressing N-terminally HA-tagged GFP fusion lacking the last β-barrel strand (HA-GFP1-10). ATOM40 and EF1a served as mitochondrial and cytosolic markers, respectively. IF analysis of cells lines allowing tetracycline-inducible expression of HA-GFP1-10 and TAC60 variants fused to the last β-strand of GFP at the C-terminus (TAC60-GFP11) or N-terminus (GFP11-TAC60), respectively. DNA is stained with DAPI (blue). ATOM40 is shown in red. GFP in green. Bar, 5 μΜ.

It has previously been shown that organellar proteins that accumulate in the cytosol upon inhibition of mitochondrial protein import are rapidly degraded by the cytosolic proteasome (28, 29). Thus to analyze whether TAC60 localizes to the outer or the inner mitochondrial membrane we followed the fate of a tagged version of the protein in inducible ATOM40- and TbTim17-RNAi cell lines. ATOM40 and TbTim17 are core subunits of the archaic protein translocase of the OM (ATOM) and the protein translocase of the IM (TIM), respectively (23, 30-34). The top panel of Fig. 4B shows that the steady state levels of tagged TAC60 in whole cells rapidly decrease during ATOM40 RNAi. The same is the case for a previously characterized β-barrel protein, the voltage dependent anion channel (VDAC), which first needs to be translocated into the intermembrane space (IMS) before it is inserted into the OM. In the TbTim17 RNAi cell line in contrast the steady state levels of TAC60 remain essentially constant. The IM protein cytochrome oxidase subunit IV (CoxIV), whose import requires TbTim17 and therefore serves as a positive control, however accumulates as unprocessed precursor protein (Fig. 4B, bottom panel). Thus mitochondrial import of tagged TAC60 depends on ATOM40 but not on TbTim17 indicating that it is an OM protein. This is further supported by a normalized abundance profile of TAC60 over six subcellular fractions, produced in a previous proteomic analysis (Fig. 4A, bottom panel) (However since TAC60 was detected in only one of two experiments, it was not included in the OM proteome defined in the study) (35, 36).

### N− and C-termini of TAC60 face the cytosol

In silico analysis of TAC60 of *T. brucei* using various prediction programs detects two high confidence TMDs (121-141 and 238-258) that are found in essentially all TAC60 orthologues of trypanosomatids. Moreover, HHPred analysis (36) indicates that the C-terminal 150 aa of TAC60 and its orthologues has some similarity to bacterial tRNA/rRNA methyltransferases.

In order to analyze the topology of TAC60 experimentally we used the split GFP approach (37). To that end a cell line expressing N-terminally HA-tagged GFP lacking the C-terminal β-strand (HA-GFP1-10) was produced. As expected the truncated GFP fractionates with the cytosol (Fig. 4C, top right panel). Subsequently, two variants of TAC60 that were N- or C-terminally fused to the last β-strand of GFP (GFP11-TAC60, TAC60-GFP11) were expressed in the same cell line. The IF analysis in Fig. 4C (bottom panel) shows a GFP signal that to a large part is in close proximity of the kDNA and thus is consistent with a TAC localization. The same signal is not seen in the absence of tetracycline, which prevents the expression of the fusion proteins. These results show that both the N- and the C-termini of TAC60 face the cytosol and thus is consistent with the notion that TAC60 has two TMDs.

### TAC60 contains distinct mitochondrial and TAC targeting signals

The integral membrane subunits of the TAC do not only need to be imported into mitochondria but also require sorting to the single unit TAC. In order to test whether mitochondrial targeting and subsequent sorting to the TAC require distinct signals, we expressed C-terminal tagged versions of TAC60 that were truncated either at their N- or C-termini or on both ends (Fig. 5A). IF analysis showed four distinct localizations of the truncated TAC60 versions (Fig. 5BCDE, Fig S1):

- The variants lacking the C-terminal 153 and 283 aa ΔC153/ΔC283) were localized to the TAC. Although some dots that do not overlap with the kDNA are also seen (Fig. 5B). The same was the case if the C-terminal 283 amino acid deletion was combined with N-terminal truncations of 75 and 97 aa (ΔN75_ΔC283/ΔN97_ΔC283) (Fig. 5B). Interestingly, however expression of these two variants causes some cells to show an enlarged kDNA or kDNA loss, respectively.
- The variant lacking the N-terminal 114 aa (ΔN114) was also localized to the TAC (Fig. 5C), although in contrast to the variants described above (Fig. 5B) the rest of the mitochondrion was also stained. Thus, only a fraction of the tagged ΔN114 variant is localized at the TAC indicating that the efficiency of TAC sorting is reduced.
- The variant lacking the N-terminal 140 aa (ΔN140) was mitochondrially localized (Fig. 5C), but not sorted to the TAC demonstrating that mitochondrial targeting and TAC-sorting are distinct events.
- The variants lacking either N-terminal 233 and 257 aa ΔN233/ΔN257) or the C-terminal 320 and 408 aa Δ320/ΔC408) finally showed a diffuse cytosolic localization (Fig. 5E).

**Figure 5.**
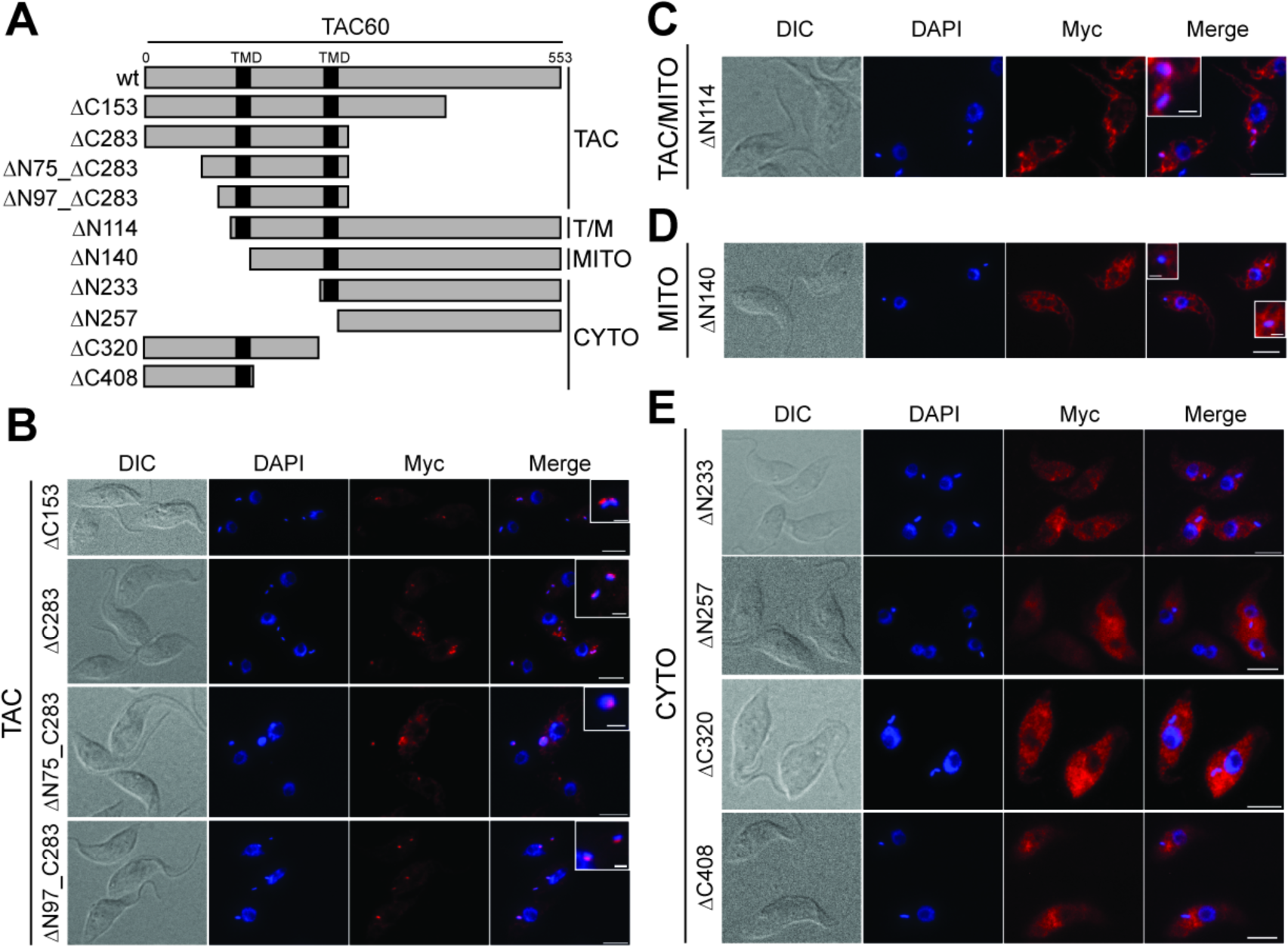
TAC60 contains distinct mitochondrial and TAC targeting signals. (A) To scale schematic drawing of the variants used to analyze TAC60 targeting. The predicted transmembrane domains of TAC60 are indicated in black. ΔNx or ΔCx indicate the number of aa deleted from the N- or/and the C-termini. All variants were C-terminally Myc-tagged. The constructs are grouped according to the four distinct localization pattern observed for the TAC60-variants: TAC, TAC and general mitochondrial (TAC/MITO or T/M), mitochondrial (MITO) and cytosolic (CYTO) localization. (B, C, D and E) IF analysis of cells lines expressing the TAC60 variants listed in (A). The cell lines are grouped as in (A). DIC, differential interference contrast picture. DNA is stained with DAPI (blue). The Myc-tagged TAC60 variants are indicated in red. The merged picture (Merge) shows an overlay of the DAPI and the Myc staining. Bar, 5 μm. Inset shows an enlargement of the kDNA region. Bar inset, μm. Co-staining with a mitochondrial marker is shown in Fig. S1.

In summary, these results define a 130 aa long segment of TAC60 - aa 140-270 - encompassing the IMS-exposed loop of the protein and the more C-terminal TMD as essential for mitochondrial targeting. Sorting to the TAC however requires an additional N-terminal 26 aa segment that consists essentially of the first TMD of TAC60.

The immunoblot in Fig. S2A confirms that all TAC60 variants are expressed in comparable amounts. However, in many cases the anti-tag antiserum detects additional signals below or above the predicted bands. The ΔN114 and ΔN140 variants show the highest heterogeneity and besides the correctly sized protein at least four major degradation products are detected as well. These degradation products are all mitochondrially loclalized (Fig. S2B) suggesting that TAC subunits that are imported into mitochondria but not sorted to the TAC are degraded. For many other variants closely spaced double bands or additional signals above the correctly sized protein are observed. Preliminary experiments indicate that at least in the case of full length TAC60 and the ΔC153 variant protein phosphatase treatment results in a more simplified pattern shifted towards a lower molecular weight range (Fig. S2C). Thus, the observed heterogeneity within the TAC60 variants might be caused by phosphorylation and possibly other postranslational modifications.

To investigate whether the correctly localized C-terminal TAC60 truncations ΔC153, ΔC283, ΔN75_ΔC283, and ΔN97_ΔC283 are functional we expressed them in a TAC60-RNAi cell line that targets part of the ORF that is absent in the truncations and therefore allows complementation experiments.

The results in Fig. 6 demonstrate that both C-terminally truncated variants complemented growth to wild-type level when expressed in the corresponding RNAi cell line. Thus the C-terminal domain of TAC60 that shows similarity to bacterial tRNA/rRNA methyltransferases is dispensable for TAC function. The same experiments were also done for the ΔN75_ΔC283 and ΔN97_ΔC283 TAC60 variants. While both of them localize to the TAC (Fig. 5B) they were not able to complement the growth phenotype indicating that their function is impaired (Fig. 6). Thus, while the N-terminal 97 aa are dispensable for TAC60 targeting they are required for the function of the protein.

**Figure 6.**
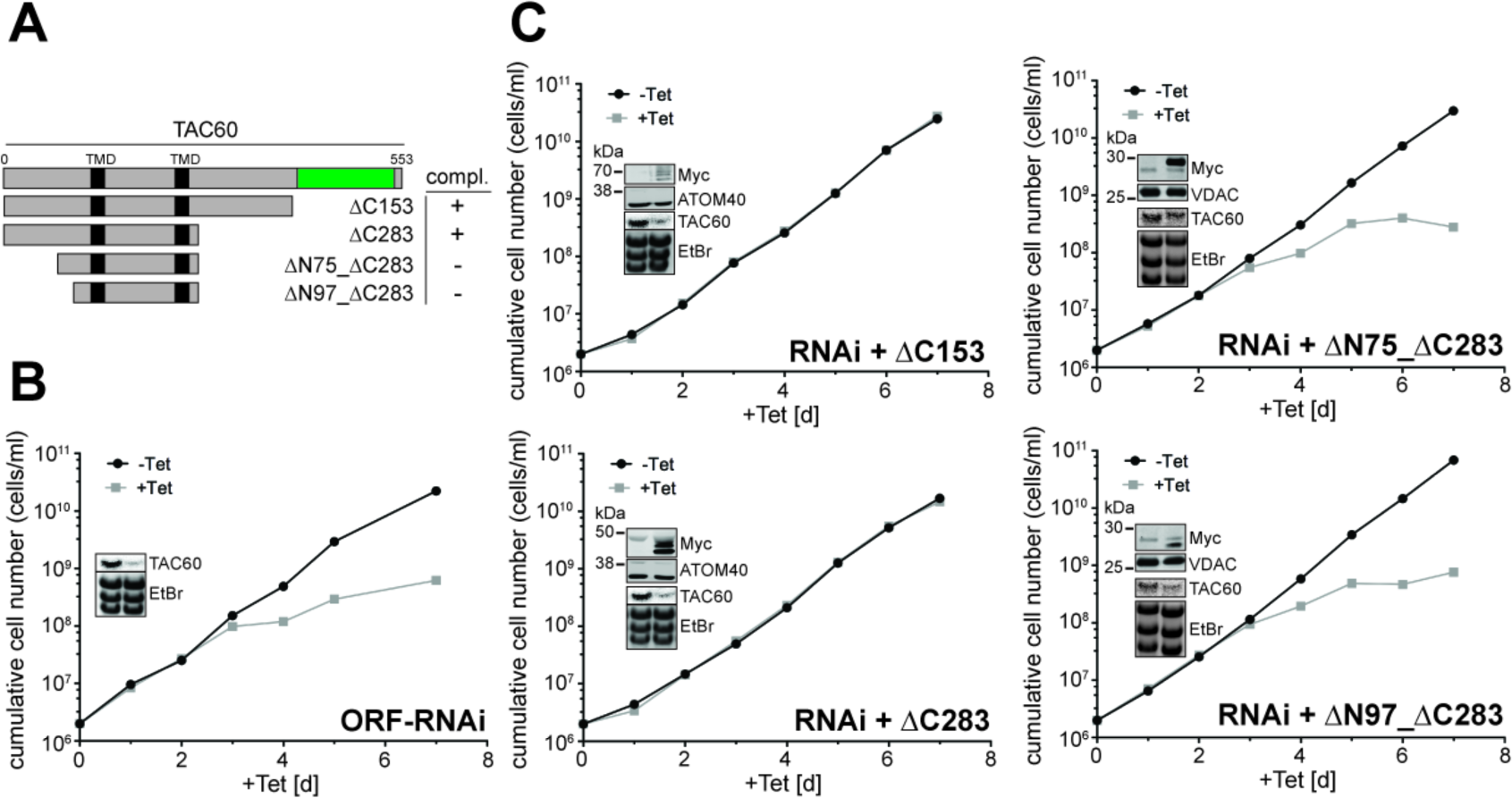
Functional analysis of TAC variants by complementation. (A) To scale schematic drawing (as in Fig. 5A) of the variants used for complementation experiments. All variants localize to the TAC. The region of the ORF that is targeted by the RNAi is shown in green. The column on the right indicates which constructs complement growth of a tetracycline-inducible TAC60-RNAi cell line targeting a sequence in the 3'-half of TAC60. (B) Growth curves of the TAC60-RNAi cell lines. The uncomplemented cell line (ORF-RNAi) and cell lines complemented with the indicated Myc-tagged versions of TAC60 (Fig. 5) are shown. Inset (top): immunoblots confirming the expression of the indicated Myc-tagged TAC60 variants. ATOM40 and VDAC serve as loading controls. Inset (bottom): Northern blots confirming the ablation of the TAC60 mRNA. Ethidiumbromide-stained (EtBr) rRNAs serve as a loading control.

### Biogenesis of TAC42 requires Sam50 and a C-terminal β-signal

Bioinformatic analysis of TAC42 does not detect any significant sequence similarity to proteins outside the kinetoplastids. As TAC60, tagged TAC42 co-fractionates with the mitochondrial marker ATOM40 when cells are extracted with low concentration of digitonin (Fig. 7A, upper panel). Moreover, the normalized abundance profile of TAC42 produced in a previous proteomic analysis suggest an OM localization (35) (as in the case of TAC60, TAC42 was only detected in one of two experiments in this study and was therefore not included in the OM proteome) (Fig. 7A, bottom panel). TAC42 is exclusively recovered in the pellet fraction in a carbonate extraction suggesting it is an integral membrane protein (Fig. 7A, middle panel). This is surprising since TAC42 lacks predicted TMDs and in silico analyses do not predict it to be a β-barrel membrane protein. Thus, to test whether the biogenesis of TAC42 depends on the β-barrel insertion machinery we expressed a C-terminally tagged version of TAC42 in a cell line allowing inducible ablation of Sam50. Interestingly, in this cell line the level of a tagged version of TAC42 decreased during RNAi (Fig. 7B). As expected the same was the case for the well characterized β-barrel proteins VDAC (38) and ATOM40 (28), which serve as positive controls. The C-terminally anchored OM protein ATOM69 (23) and cytosolic translation elongation factor 1a (EF1a) on the other hand were not affected.

**Figure 7.**
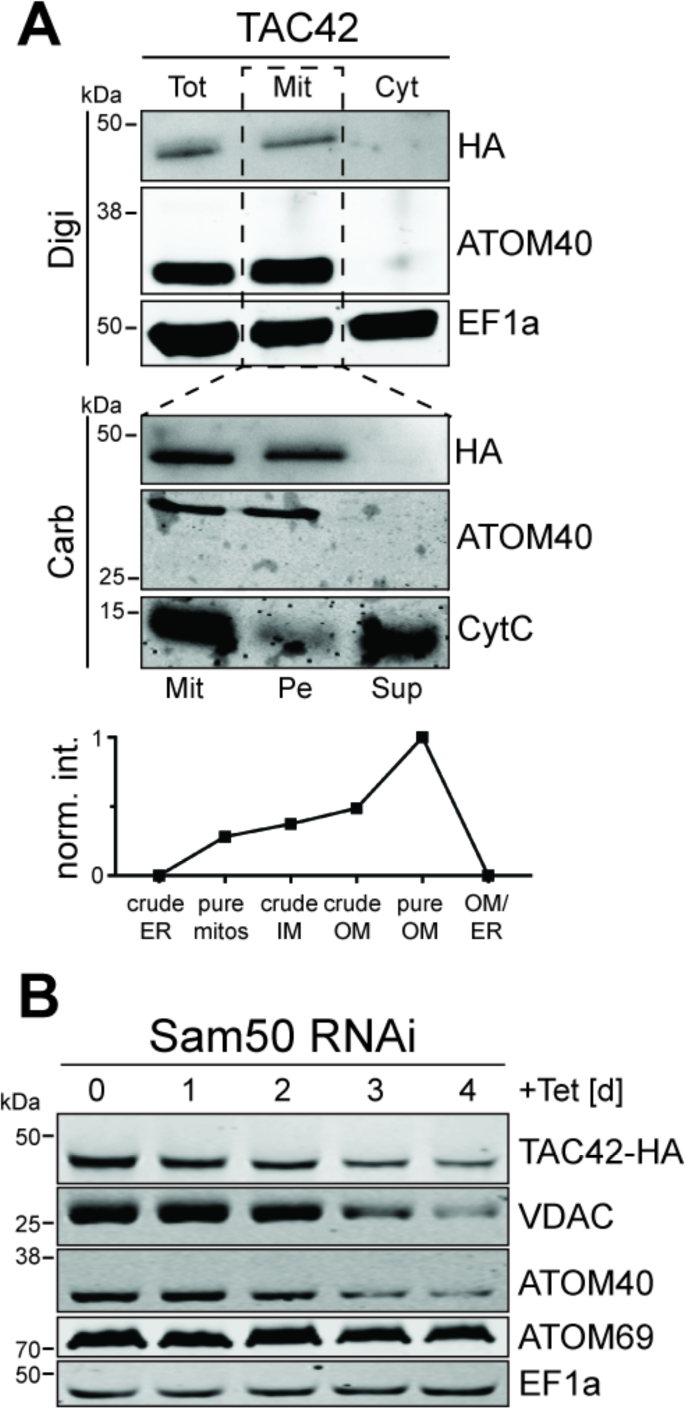
TAC42 is an OM protein whose biogenesis depends on Sam50. (A) Top panel: immunoblot analysis of whole cells (Tot), digitonin-extracted mitochondria-enriched pellet (Mit) and soluble (Cyt) fractions of cells expressing C-terminally HA-tagged TAC42. ATOM40 and EF1a served as mitochondrial and cytosolic markers, respectively. Middle panel: carbonate extraction at ph 11.5 of the mitochondria-enriched pellet fraction (Mit). The pellet (Pe) and the supernatant (Sup) fractions correspond integral membrane and soluble proteins, respectively. ATOM40 and Cyt C serve as markers for integral and peripheral membrane proteins, respectively. Bottom panel: normalized abundance profile of TAC42 over six subcellular fractions, produced in a previous proteomic analysis (35). (B) Immunoblots of total cellular extracts from procyclic Sam50-RNAi cells that constitutively express HA-tagged TAC42. Time of induction is indicated. The mitochondrial β-barrel proteins VDAC and ATOM40, and the α-helically anchored OM protein ATOM69 serve as positive and negative controls, respectively. Cytosolic EF1a was used as a loading control.

A pioneering study in yeast identified a moderately conserved sequence in the last β-strand of mitochondrial β-barrel proteins that serves as a sorting signal which directs the protein to the β-barrel protein insertion machinery (39). Fig. 8A shows that the three trypanosomal β-barrel proteins TAC40, ATOM40 and VDAC as well as TAC42 have C-termini corresponding to the β-signal consensus sequence. Thus, in order to test whether these sequences function as sorting signals we expressed tagged TAC42 and TAC40 variants containing mutated variants of the putative β-signals. In one variant, termed 1mut, the invariant glycine was mutated to alanine. In the other variant, termed 4mut, all four conserved positions were changed, the first two to alanines and the last two to serines.

**Figure 8.**
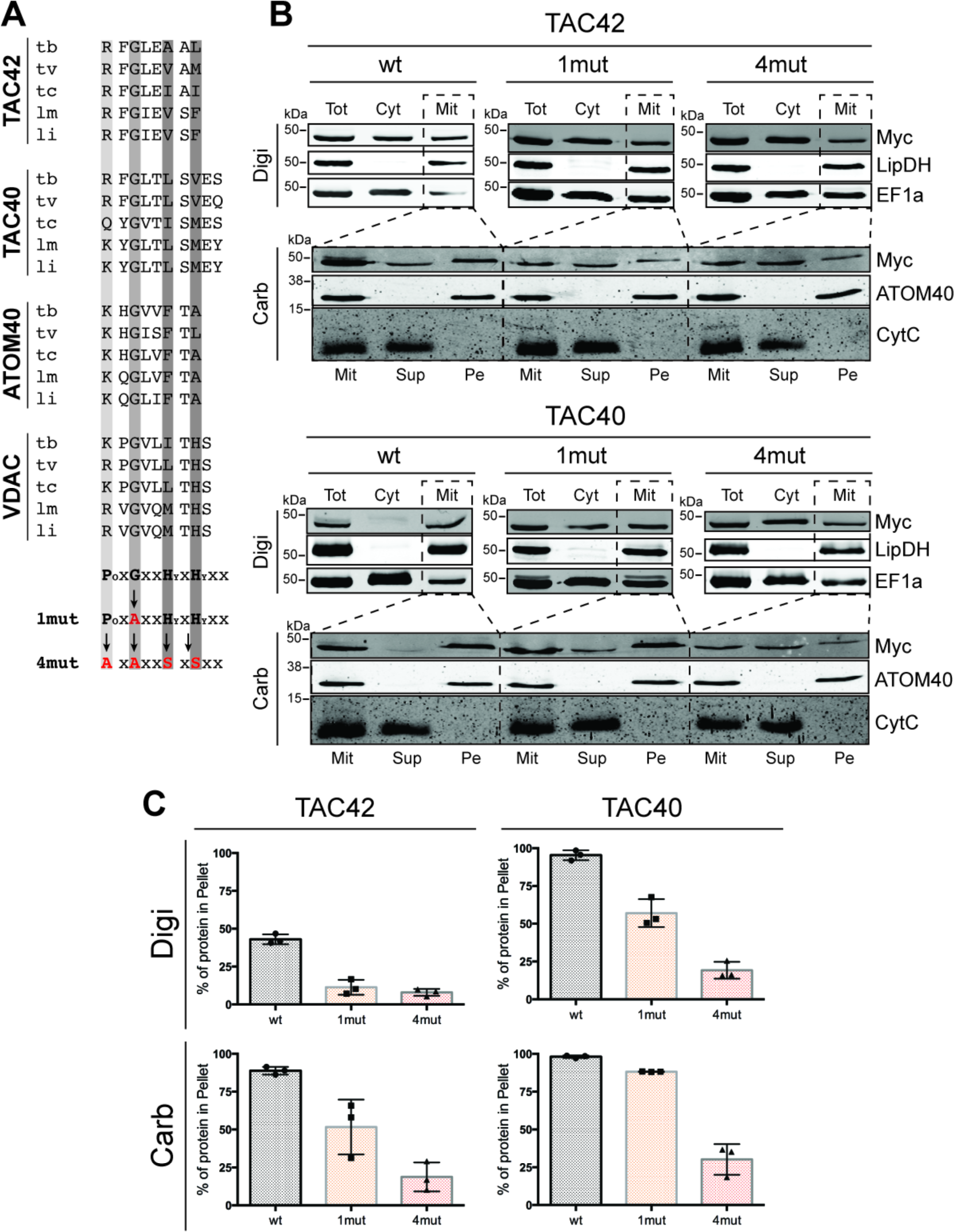
TAC42 and TAC40 contain C-terminal β-signals. (A) Amino acid sequence of the C-termini of the indicated β-barrel proteins in different trypanosomatids. Tb, *T. brucei;* T. v, *T. vivax;* Tc, *T. cruzi;* Lm, *L. major* and Li, *L. infantum*. The residues contributing to the predicted β-signal are indicated by the grey bars. The β-signal consenus sequence and the sequence of the two mutated variants (1mut, 4mut) of the β-signal consensus sequence are indicated. Changed residues are shown in red. Po, polar amino acid; Hy, hydrophobic amino acid. (B) TAC42 top panel: immunoblot analysis of whole cells (Tot), soluble (Cyt) and digitonin-extracted mitochondria-enriched pellet (Mit) fractions of cell lines expressing C-terminally Myc-tagged full length (wt), 1mut and 4mut variants of TAC42, respectively. Lipoamide dehydrogenase (LipDH) and EF1a serve as mitochondrial and cytosolic markers, respectively. TAC42 bottom panel: carbonate extraction at pH 11.5 of the mitochondria-enriched pellet fraction (Mit) of the cell lines depicted above. The pellet (Pe) and the supernatant (Sup) fractions correspond integral membrane and soluble proteins, respectively. AT0M40 and Cyt C serve as markers for integral and peripheral membrane proteins, respectively. TAC40 top and bottom panels, same experiments as for TAC42 and its variants were performed for TAC40 and its two mutated variants. (C) Graphs showing the mean and the standard errors of a quantification of three biological replicates of the experiment shown in (B).

The digitonin extraction in the top panel of Fig. 8B shows that approximately 45% of the tagged wildtype version of TAC42 is recovered in the pellet corresponding to a crude mitochondrial fraction. The fact that 55% of the tagged proteins remains in the supernatant is likely due to overexpression when compared to the endogenous protein. Interestingly, however in the case of the 1mut and 4mut versions of TAC42 only 8-11% of the proteins are recovered in the pellet fractions. The same experiments were also done with the previously characterized β-barrel protein TAC40 (22) and, as in the case of TAC42, mitochondrial targeting of the 1mut and 4mut versions of TAC40 was dramatically impaired (Fig. 8B). TAC40 was also ectopically tagged but likely less overexpressed compared to TAC42 which may explain why almost all of the tagged wildtype variant of the protein is mitochondrially localized. Thus, mutating the conserved glycine or all conserved amino acids in the β-signal consensus sequence of TAC42 and TAC40 progressively reduces mitochondrial targeting of the two proteins.

The β-signal mediates the interaction with the β-barrel insertion machinery (39). Its absence is therefore expected to interfere with membrane insertion after the proteins have been translocated into the IMS. In order to test this prediction we analyzed the mitochondria-associated fractions of the tagged TAC42 and TAC40 and its corresponding 1mut and 4mut versions by carbonate extraction at high pH to determine whether the proteins have been inserted into the OM. The lower panels in Fig. 8B show that the tagged wildtype TAC42 and TAC40 are recovered in the pellet fractions together with ATOM40, which serves as marker for a correctly inserted integral membrane protein. However, membrane insertion of the 1mut variant of TAC42 is reduced by 50% and in the case of 4mut variants by 70-80% for both proteins. Thus, TAC42 and TAC40 contain a C-terminal β-signal that is essential for correct targeting and membrane insertion of the proteins into the mitochondrial OM.

The β-signal is expected to be conserved in all eukaryotes (39). In order to show direct interaction of the putative β-barrel protein TAC42 with the SAM complex we therefore performed *in vitro* import experiments using trypanosomal substrate proteins and isolated yeast mitochondria. Radioactive substrates, produced by in vitro translation using rabbit reticulocyte lysate, were incubated with isolated mitochondria from either wildtype *S. cerevisiae* or from a yeast strain carrying a deletion of the SAM complex subunit Sam37, whose function is to promote β-barrel protein insertion into the OM by linking the SAM and the TOM complex (40, 41). The *in vitro* import reactions were analyzed by BN-PAGE. The results in Fig. 9A show that the wild-type protein but not the 4mut variant of TAC42 accumulate in a time dependent manner in a complex of approximately 200 kDa. Moreover, when the wild-type TAC42 is imported into mitochondria lacking Sam37 a much smaller amount of the complex is observed. Furthermore, its molecular weight is slightly lower due to the absence of Sam37 (42). In summary these results show that trypanosomal TAC42 directly interacts with the yeast SAM complex in a β-signal and Sam37-dependent manner indicating that TAC42 is indeed a novel kinetoplastid-specific mitochondrial β-barrel protein.

**Figure 9.**
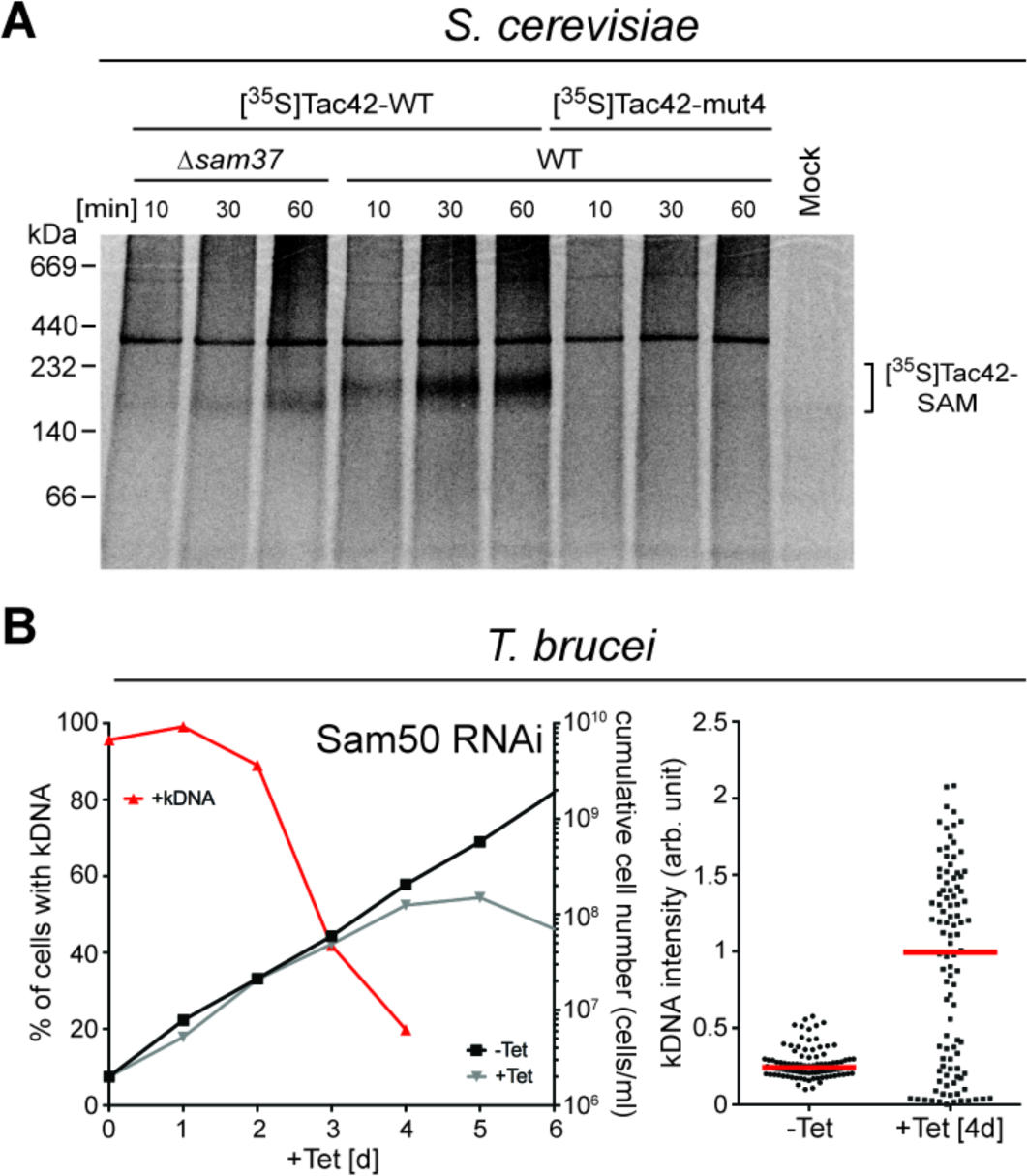
Trypanosomal β-signals are functional in yeast and knock-down of trypanosomal Sam50 causes a kDNA segregation defect. (A) In vitro translated ^35^S-labeled TAC42 and TAC42-mut4 were incubated for the indicated time with mitochondria either isolated from wildtype (WT) yeast or from a yeast strain lacking the SAM complex subunit sam37 (*Asam37*). Subsequently all samples were analyzed by BN-PAGE and analyzed by autoradiography. A mock reaction lacking organelles was also analyzed. The putative TAC42/Sam50 complex is indicated. (B) Left graph: growth and loss of kDNA in the procyclic *T. brucei* Sam50-RNAi cell line. Red lines depict percentage of cells still having the kDNA. Right graph: fluorescent intensities of kDNA networks were measured after Sam50 knock down. Red lines mark the median.

Thus at least two essential OM subunits of the TAC - TAC40 and TAC42 - are β-barrel proteins. In line with this, ablation of Sam50, the core component of the SAM complex, leads to a rapid increase of cells lacking kDNA networks, whereas in the cells that have retained the kDNA it is massively overreplicated (Fig. 9B). Ablation of Sam50 therefore essentially reproduces the phenotypes that are observed after ablation of individual TAC subunits. However, the same is not seen in cells ablated for ATOM40 the channel subunit of the main OM protein translocase (20). Thus these results underscore the importance of the trypanosomal SAM complex for the assembly of this unique mitochondrial DNA inheritance system.

## Discussion

The TAC is a single unit structure that links the kDNA to the basal body. During each cell cycle a new TAC needs to be formed to guarantee that the replicated kDNA networks are correctly segregated during the binary fission of the single mitochondrion of *T. brucei*. Understanding TAC biogenesis is hampered by the fact that only few of its subunits are known and that their targeting pathways have not been studied. The exclusion zone filaments, that form the bridge from the basal body to the OM, consists of cytosolic proteins which may reach the TAC region by diffusion and therefore not require specific targeting. The subunits of the unilateral filaments, however, need to be imported into the mitochondrial matrix before they can assemble to link the kDNA network with the mitochondrial IM. A similar situation is found for the membrane-embedded TAC subunits in the differentiated membranes which connect the cytosolic with the matrix-localized TAC filaments. Its subunits potentially first need to be imported and inserted into the OM and IM membranes, respectively, before they are laterally sorted to the new TAC that is being assembled.

In our studies we have focused on the OM region of the differentiated membrane domain of the TAC. We have discovered two novel integral OM TAC subunits - TAC60 and TAC42, that are required for kDNA replication and segregation - determined their topology and deciphered their biogenesis pathways.

TAC60 is essential for TAC function, contains two TMDs and its N- and C-termini both face the cytosol. In a deletion analysis we have uncoupled mitochondrial targeting from TAC sorting and identified the first TAC sorting signal. Mitochondrial targeting of TAC60 requires a 143 long region that includes the IMS-exposed loop and its more C-terminal TMD. Insertion of TAC60 into the mitochondrial OM is mediated by ATOM40, the pore-forming subunit of the master protein translocase in the OM. However, to reach its final destination, the TAC, TAC60 requires an additional 26 aa comprising the first TMD. Whether the TAC sorting signal depends on the TAC60 mitochondrial targeting signal or whether it could in principle work on its own, provided that the first TMD of TAC60 is inserted into the OM in the correct topology, remains unknown at the moment. Moreover, presently the TAC sorting signal appears to be specific for TAC60 indicating that other TAC subunits may have different sorting signals.

The essential TAC subunit TAC42 lacks predictable TMDs. Its localization to the TAC depends on both Sam50, the core component of the SAM complex, and on the presence of a β-signal consensus sequence at its C-terminus. It has been shown in yeast that this sequence mediates the interaction of β-barrel proteins with the SAM complex (39). Moreover, TAC42 can be inserted into the OM of isolated yeast mitochondria provided that they have a functional SAM complex and that it carries a functional β-signal. In summary these result demonstrate that TAC42 is mitochondrial β-barrel protein even though in silico analysis fails to predict so.

*T. brucei* has six known β-barrel membrane proteins: the metabolite transporter VDAC (38), a second VDAC-like protein of unknown function (43), ATOM40 (28, 44) and Sam50 (45), the core components of the ATOM and the SAM complex, and finally TAC40 (22) and TAC42 essential subunits of the TAC. Four of them (VDAC, the VDAC-like protein, ATOM40, TAC40) belong to the mitochondrial porin protein family of whereas Sam50 is an Omp85-like protein. TAC42 is unique it is neither a mitochondrial porin nor an Omp85-like protein but defines a novel class kinetoplastid-specific β-barrel proteins essential for mitochondrial DNA inheritance.

The presence of two distinct β-barrel proteins, TAC40 and TAC42, in the TAC is striking. β-barrel proteins are exclusively found in the OMs of bacteria, mitochondria and plastids (41, 46). It can be speculated that their prominent presence in the TAC indicates that the ancestor of this DNA inheritance system evolved very early, at a time when the integral membrane proteins present in the OM of the mitochondrial ancestor were restricted to β-barrel proteins.

Moreover, the membrane domain of the TAC shows some architectural similarity to the double membrane spanning secretion systems of gram negative bacteria (47). Both types of structures link OM and IM of bacterial evolutionary origin. Moreover, the OM is in both cases spanned by a β-barrel type structure (generally multimeric in the case of bacteria) and thus requires a Sam50/BamA-type insertion system. However, whereas the bacterial secretion systems serve to export bacterial effector proteins, there is no evidence that the TAC is involved in transport processes.

While many subunits of the TAC are still unknown we get a more and more detailed picture of its OM constituents. Up to now four essential, integral OM subunits of the TAC have been characterized: the β-barrel proteins TAC40 and TAC42, which form a complex with TAC60, as well as the dually localized pATOM36. This complexity is surprising since the TAC is expected to have a structural function, linking the kDNA network to the basal body of the flagellum. A single OM protein that interacts on the cytosolic side with the exclusion zone filaments and on the IMS side with an IM protein should in principle be sufficient to do this job. We would like to propose two possible explanations for this unexpected complexity.

It could be that the function of the TAC goes far beyond providing a structural linkage. Being localized between the kDNA and the flagellum, the TAC would be ideally suited to serve as a signaling platform that for example may regulate and integrate kDNA replication and segregation with flagellar growth and cytokinesis. Indeed pATOM36 has already been shown to mediate both mitochondrial protein import and mitochondrial DNA inheritance (20).

Alternatively, it might be that the TAC is the product of constructive neutral evolution. This ratched-like evolutionary process provides a non-adaptive explanation why macromolecular complexes can be comprised of more subunits than their function seem to demand (48, 49). In the case of the TAC, a possible scenario would be that an autonomously functioning ancestral TAC subunit would fortuitously bind to another protein. Binding to this protein would not affect the function of the TAC subunit, but it would have the potential to suppress mutations, which if present in the absence of the binding partner would inactivate the TAC subunit. Should such mutations occur the TAC subunit would lose its autonomy, as its function would now depend on the other protein. Thus, constructive neutral evolution may have led to four or more OM TAC subunits even though common sense suggests a single one should be enough.

The two proposed explanations are not mutually exclusive, as both may have contributed to the complex TAC architecture. In order to disentangle the two we need a more complete picture of the TAC composition and architecture as well as a detailed functional analysis of its subunits.

## Materials and Methods

### Transgenic cell lines

Transgenic procyclic cell lines are based on *T. brucei* 29-13 (50) and were cultured at 27°C in SDM-79 containing 10% (v/v) fetal calf serum (FCS). Transgenic bloodstream form trypanosomes are based on the New York single marker (NYsm) strain or on a derivative thereof termed F_1_γL262P (27). All bloodstream from cells were grown at 37°C in HMI-9 supplemented with 10% FCS (v/v).

Full length TAC60 (Tb927.7.1400) (Fig. 1B, Fig. 2A) or deletion variants of it (Fig. 5, Fig. 6, Fig. S1 and S2), termed ΔN114 (nt 343-1662), ΔN140 (nt 421-1662), ΔN233 (nt 697-1662), ΔN257 (nt 772-1662), ΔN153 (nt 1-1200), ΔC283 (nt 1-810), ΔN75_ΔC283 (nt 226-810), ΔN97_ΔC283 (nt 292-810), ΔC320 (nt 1-696), ΔC408 (nt 1-435) were cloned into a modified pLew100 expression vector containing a puromycin resistance gene in which a cassette had been inserted allowing C-terminal triple Myc-tagging (51).

For the experiment shown in Fig. 4A, B one allele of TAC60 was tagged in situ at the C-terminus with a triple c-Myc-epitope (51) in the background of procyclic RNAi cell lines targeting ATOM40 and TbTim17, both of which have been described before (28, 52).

Full length TAC42 (Tb927.7.3060) (Fig. 8) and TAC40 (Fig. 8) or mutated variants of them (Fig. 8), termed 1mut (TAC40: G351A; TAC42: G382A) or 4mut (TAC40: R349A, G351A, L354S and V356S; TAC42: R380A, G382A, A385S and L387S) were cloned into a modified pLew100 expression vector containing a puromycin resistance gene in which a cassette had been inserted allowing C-terminal triple Myc-tagging (51). For the experiment shown in Fig. 3A and Fig. 7 one allele of TAC42 was tagged in situ at the C-terminus with a triple HA-epitope and expressed in *T. brucei* 29-13 cells and in a previously described RNAi cell line targeting Sam50 (52) [5]. The in situ HA-tagged TAC40 cell line (Fig. 1A) has been described before (22). The RNAi in the procyclic and bloodstream form cell lines was targeted against the ORF of TAC60 (nt 256-772) or TAC42 (nt 238-710), respectively. For the complementation experiments of TAC60 (Fig. 6), a different RNAi cell line targeting the 3'-part of the TAC60 ORF (nt 1220-1629) was established.

The Split-GFP approach was used as described (53). For the results shown in Fig. 4C, the GFP 1-10 OPT (GFP1-10) and the M3 strand 11 (GFP11) were amplified from a pET-15b-based vector (generous gift from Prof. Steven Boxer, University of Stanford). GFP 1-10 OPT was cloned into a pLew100-based expression vector containing a blasticidin resistance cassette. The resulting construct allows inducible expression of N-terminal HA-tagged cytosolic GFP1-10 (HA-GFP1-10). GFP11 was cloned into a pLew100 expression vector containing the puromycin resistance gene. The resulting construct allows C-terminal tagging of proteins with GFP11. Subsequently, the complete ORF of TAC60 was cloned into this modified pLew100 vector, yielding the construct TAC60-GFP11. Another pLew100 expression vector was established, which allows N-terminal tagging of proteins with GFP11. Subsequently, the complete ORF of TAC60 was cloned into this modified pLew100 vector, yielding the construct GFP11-TAC60. A *T. brucei* 29-13 cell line expressing HA-GFP1-10 was established, which subsequently was transfected with TAC60-GFP11 or GFP11-TAC60, respectively.

For visualization of tagged proteins, the respective cell lines were induced for 1 day with 1 μg/ml of tetracycline.

### Antibodies

The following polyclonal rabbit antisera directed against the indicated antigens were produced in our lab and have been used before (23, 35). The working dilutions for immunoblots (IB) and IF are indicated: VDAC (IB 1:1,000), ATOM40 (IB 1:10,000; IF 1:1,000), cytochrome C (IB 1:1,000), ATOM69 (IB 1:50), LipDH (IB 1:10,000) and CoxIV (IB 1:1,000). Commercially available monoclonal antibodies were used as follows: mouse c-Myc (Invitrogen, 132500; IB 1:2,000; IF 1:50), mouse HA (Enzo Life Sciences AG, CO-MMS-101 R-1000; IB 1:5,000; IF 1:1,000) and mouse EF1a (Merck Millipore, Product No. 05-235; IB 1:10,000). Monoclonal anti-tyrosinated α-tubulin antibody YL1/2 (54) (IFA 1:500) produced in rat was a generous gift from Prof. Keith Gull, University of Oxford.

Secondary antibodies for IB analysis were IRDye 680LT goat anti-mouse, IRDye 800CW goat anti-rabbit (LI-COR Biosciences, 1:20,000) and horse radish peroxidase-coupled goat anti-mouse and anti-rabbit (Sigma-Aldrich, 1:5,000). Secondary antibodies for IF were goat anti-mouse Alexa Fluor 633, goat anti-mouse Alexa Fluor 596, goat anti-rabbit Alexa Fluor 488 and goat anti-rat Alexa Fluor 488 (all from ThermoFisher Scientific, 1:1000)

### Digitonin extraction

Digitonin extraction was used to generate crude mitochondrial enriched fractions (55) to demonstrate mitochondrial localization of a protein of interest. For this, 5x10^7^ or 1x10^8^ cells were incubated for 10 min on ice in 20 mM Tris-HCl pH 7.5, 0.6 M sorbitol, 2 mM EDTA containing 0.025 % (w/v) digitonin. After centrifugation (6,800 g, 4°C), the resulting mitochondria-enriched pellet was separated from the supernatant and equal cell equivalents of each fraction were subjected to SDS-PAGE and immunoblotting. Alternatively, the mitochondria-enriched pellets were used for subsequent alkaline carbonate extractions (see below).

### Alkaline carbonate extraction

A mitochondria-enriched pellet fraction obtained by digitonin extraction was resuspended in 100 mM Na2CO3 pH 11.5, incubated on ice for 10 min and centrifuged (100,000 g, 4°C, 10 min) to separate the membrane fraction from soluble proteins. Equal cell equivalents of all samples were analyzed by SDS-PAGE und immunoblotting.

### SILAC proteomics and immunoprecipitations

SILAC-IP experiments were essentially done as described (20). *T. brucei* 29-13 cells, their derivatives allowing expression of in situ HA-tagged TAC40 or TAC42, and a transgenic cell line allowing inducible expression of Myc-tagged TAC60 were used as indicated (Fig. 1). Cells expressing or not expressing the tagged bait protein were grown in either light (unlabeled) or heavy (^13^C6^15^N4-L-arginine; ^13^C6^15^N2-lysine) arginine- and lysine-containing SDM-80 medium supplemented containing 10-15% (v/v) dialyzed FCS (BioConcept, Switzerland) for around 10 doubling times to establish complete labeling of proteins with light or heavy amino acids.

Equal numbers of cells grown in the presence of heavy or light arginine and lysine were mixed and harvested. The resulting pellets were solubilized in 20 mM Tris-HCl, pH 7.4, 0.1 mM EDTA, 100 mM NaCl, 10% glycerol, 1.5% (w/v) digitonin and 1X Protease Inhibitor mix (EDTA-free, Roche) for 15 min at 4°C. The extracts were centrifuged (20,000 g, 15 min, 4°C) and the resulting supernatants were incubated with anti-HA affinity beads (Roche) or anti-Myc affinity beads (EZview red, Sigma), equilibrated in the same buffer as above but containing only 0.2% (w/v) of digitonin. After 2 h of incubation at 4°C, the supernatant was discarded and the beads were washed 3 times with 0.5 ml of the same buffer. TAC40 and TAC60 protein complexes were eluted by boiling the resin for 5 min in 60 mM Tris-HCl, pH 6.8 containing 0.1% SDS whereas TAC42 protein complexes were eluted using SDS gel loading buffer without β-mercaptoethanol. SILAC-IP experiments were performed in three biological replicates including a label-switch each.

### Quantitative liquid chromatography-mass spectrometry ( LC-MS)

Proteins purified in TAC42 SILAC-IPs were separated on a 4-20% Mini Protean TGX SDS-PAGE gel (BioRad). Afterwards, the gel was fixed with acetic acid and methanol and stained for 3 hours with fresh ammonium sulfate-based colloidal coomassie (10% phosphoric acid, 10% ammonium sulfate, 0.12% coomassie brilliant blue G, 20% methanol). Subsequently, the gel lanes were cut into 10 pieces per replicate, which were then destained by repetitive rounds of incubation in 10 mM NH4HCO3 and 10 mM NH4HCO3 containing 50% ethanol. When completely destained, the gel pieces were dehydrated in 100% ethanol. Finally, cysteine residues of proteins were reduced with 5 mM bond breaker solution (Thermo Fisher Scientific), alkylated with 100 mM iodoacetamide in 10 mM NH4HCO3, and proteolytically digested with trypsin (37°C, incubation overnight). Proteins purified in TAC40 and TAC60 SILAC-IPs were were reduced, alkylated and tryptically digested in-solution as described previously (29).

LC-MS analyses of peptide mixtures were performed on an LTQ Oritrap XL (TAC40, TAC60) or an Orbitrap Elite (TAC62) mass spectrometer (Thermo Fisher Scientific, Bremen, Germany), each directly coupled to an UltiMate 3000 RSLCnano HPLC system (Thermo Fisher Scientific, Dreieich, Germany), as described before (32). For quantitative MS data analysis, MaxQuant/Andromda was used (version 1.4.1.2 for TAC40, 1.5.1.0 for TAC60, and 1.5.3.30 for TAC42 data; (56, 57)). MS/MS data were searched against all entries for *T. brucei*TREU927 retrieved from the TriTryp database (version 8.1) applying MaxQuant default parameters with the exceptions that protein identification and quantification were based on one unique peptide and one ratio count (i.e. SILAC peptide pair). Mean log10 protein abundance ratios (TAC40/42/60 versus control) and p-value (one-sided Student's t-test) across at least two 2 biological replicates were determined. Proteins identified and quantified in TAC40, TAC42, and TAC60 SILAC IPs are listed in Tables S1 - S3.

### In vitro import and assembly assays using mitochondria from *S. cerevisiae*

^35^S-Met-labelled proteins were synthesized using the TNT T7 Quick for PCR (Promega) in vitro translation kit according to the instruction manual. Radiolabeled precursors proteins were incubated with isolated yeast mitochondria in import buffer (3% (w/v) BSA, 250 mM sucrose, 80 mM KCl, 5 mM MgCl2, 5 mM L-methionine, 2 mM KH2PO4, 10 mM MOPS-KOH, pH 7.2, 2 mM NADH, 5 mM ATP, 10 mM creatine phosphate, 0.1 mg/ml creatine kinase) at 25°C. Mitochondria were washed with SEM (250 mM sucrose, 1mM EDTA, 10mM MOPS pH 7.2) and analyzed by blue native electrophoresis (4-16% gradient gels) and autoradiography.

### Miscellaneous

IF and Northern blots were done as described (23). IF images were acquired with a DFC360 FX monochrome camera (Leica Microsystrems) and a DMI6000B microscope (Leica Microsystems). Image analysis was done using LAS X software (Leica Microsystems), ImageJ, and Adobe Photoshop CS5.1 (Adobe). Relative quantification of the fluorescent intensity of kDNA networks is described in (22). Isolation of flagella was done according to (58).

## Acknowledgments

We thank Elke Horn and Bettina Knapp for technical assistance. Research in the groups of B. W. and C.M. was funded by the Deutsche Forschungsgemeinschaft and the Excellence Initiative of the German Federal & State Governments (EXC 294 BIOSS Centre for Biological Signalling Studies) and the RTG2202. Research in the lab of A. Schneider was supported by grant 138355 and in part by the NCCR "RNA & Disease" both funded by the Swiss National Science Foundation.

---

**Figure S1.**
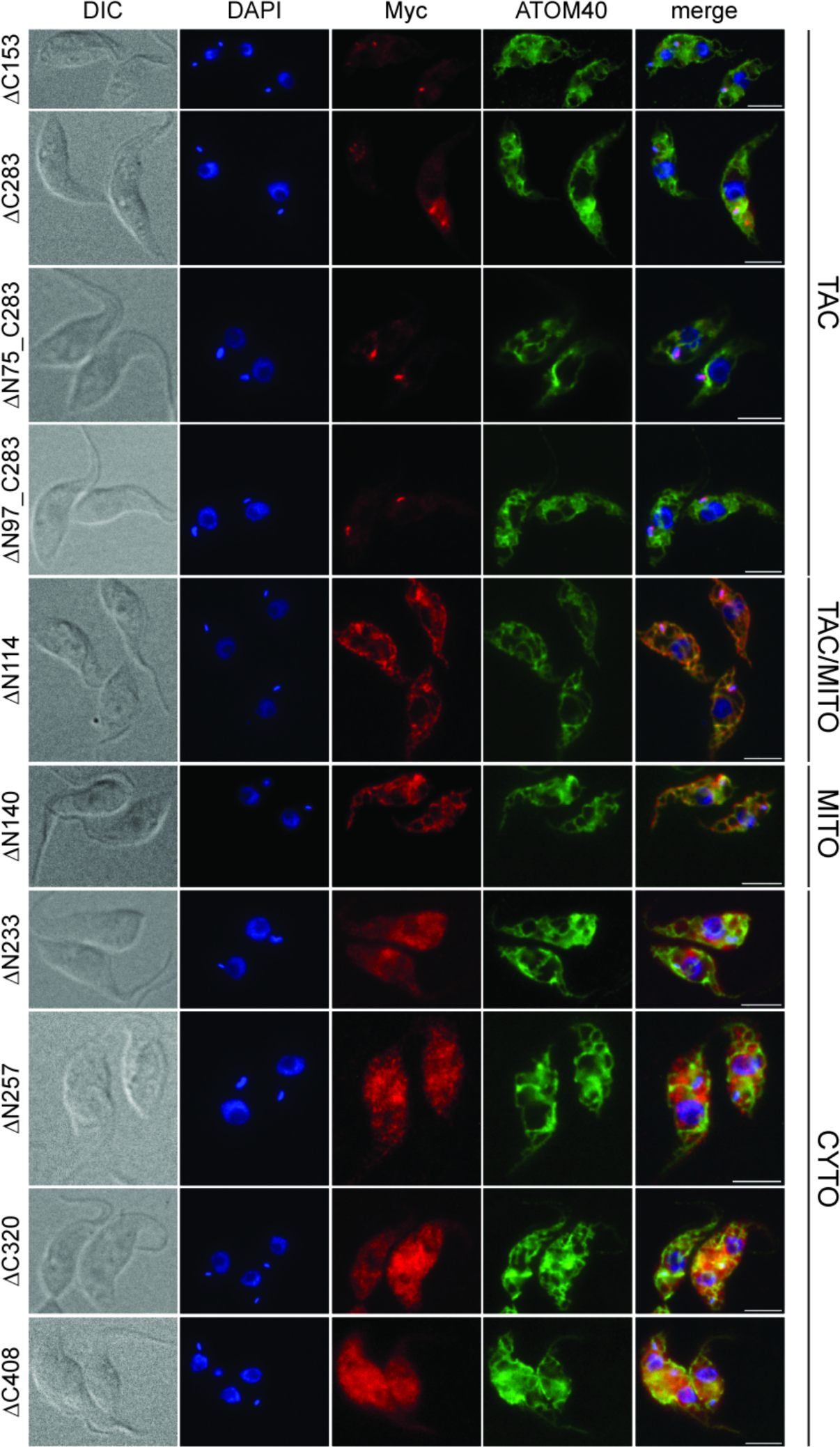
IF analysis of cell lines expressing TAC60 variants. The cell lines are grouped according to the localization pattern of the TAC60-variants as in shown in Fig. 5. DIC, differential interference contrast picture. DNA is stained with DAPI (blue). The Myc-tagged TAC60 variants are indicated in red. The mitochondrial marker AT0M40 is shown in green. The last column shows the merge of all three signals. Co-localization of the TAC60-variants with the kDNA results in a purple signal. Co-localization of the TAC-variants with the mitochondrial marker results in a yellow staining. Bar, 5 μm.

**Figure S2.**
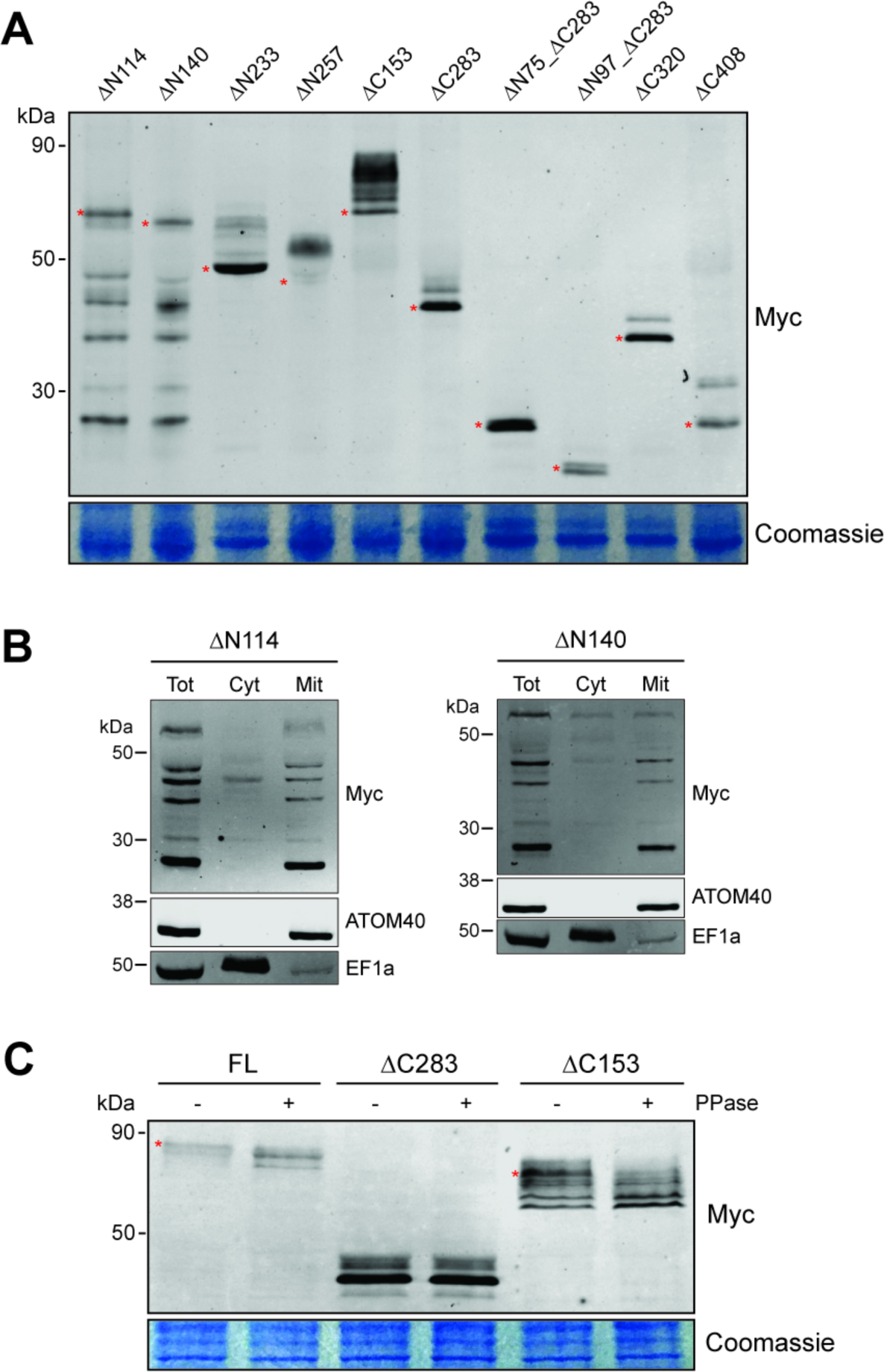
Expression of TAC60 variants. (A) Immunoblot of an SDS-PAGE containing total cellular extracts of the indicated Myc-tagged TAC60-variant expressing cells. Red asterisks indicate which bands of the TAC60-variants most closely match their calculated molecular weight. (B) Immunoblot analysis of whole cells (Tot), soluble (Cyt) and digitonin-extracted mitochondria-enriched pellet (Mit) fractions of cells expressing either the C-terminally Myc-tagged ΔN114 (left panel) or ΔN140 (right panel) TAC60 variant. AT0M40 and EF1a served as mitochondrial and cytosolic markers, respectively. (C) Protein phosphatase (PPase) treatment of total cellular extracts derived from cells expressing the indicated constructs suggests that full length TAC60 and the ΔC153 variant are phosphorylated. Red asterisks indicate which bands are affected by the PPase treatment. The bottom panels in (A) and (C) show a section of the corresponding Coomassie-stained gels that serve as loading controls.

